# Integrating Comprehensive Functional Annotations to Boost Power and Accuracy in Gene-Based Association Analysis

**DOI:** 10.1101/732404

**Authors:** Corbin Quick, Xiaoquan Wen, Gonçalo Abecasis, Michael Boehnke, Hyun Min Kang

**Affiliations:** Department of Biostatistics and Center for Statistical Genetics, University of Michigan, Ann Arbor; Department of Biostatistics, Harvard T. H. Chan School of Public Health; Regeneron Pharmaceuticals

## Abstract

Gene-based association tests aggregate genotypes across multiple variants for each gene, providing an interpretable gene-level analysis framework for genome-wide association studies (GWAS). Early gene-based test applications often focused on rare coding variants; a more recent wave of gene-based methods, e.g. TWAS, use eQTLs to interrogate regulatory associations. Regulatory variants are expected to be particularly valuable for gene-based analysis, since most GWAS associations to date are non-coding. However, identifying causal genes from regulatory associations remains challenging and contentious. Here, we present a statistical framework and computational tool to integrate heterogeneous annotations with GWAS summary statistics for gene-based analysis, applied with comprehensive coding and tissue-specific regulatory annotations. We compare power and accuracy identifying causal genes across single-annotation, omnibus, and annotation-agnostic gene-based tests in simulation studies and an analysis of 128 traits from the UK Biobank, and find that incorporating heterogeneous annotations in gene-based association analysis increases power and performance identifying causal genes.

## Introduction

Genome-wide association studies (GWAS) have identified thousands of genetic loci associated with complex traits (Welter et al. 2013); however, the biological mechanisms underlying these associations are often poorly understood. Gene-based association tests can provide a more interpretable analysis framework compared to single-variant analysis, interrogating association at the gene level by aggregating genotypes across multiple variants for each gene. This strategy can also increase power to detect association by aggregating small effects across variants, reducing the burden of multiple testing, and weighting or filtering to prioritize functional variants (Neale and Sham 2004; Sham and Purcell 2014).

In gene-based analysis, variants are often grouped or weighted by putative functional effect, for example, a common strategy for exome analysis is to include only rare non-synonymous or loss-of-function (LoF) variants in gene-based tests such as SKAT and the CMC burden test (DJ Liu et al. 2014; Morrison et al. 2013). A more recent wave of gene-based methods, e.g. PrediXcan (Gamazon et al. 2015; A Barbeira et al. 2016) and TWAS (Gusev et al. 2016), use eQTL variants (eVariants) to construct gene-based tests of association between the predicted genetic component of gene expression and GWAS trait. Incorporating regulatory variants is expected to be particularly valuable for gene-based analysis of complex traits, since most genetic associations discovered to date are in non-coding regions (MacArthur et al. 2016). However, while coding variants generally implicate a single known gene, the gene(s) affected by regulatory variants are often less clear (Ernst et al. 2011; Cao et al. 2017).

Incorporating multiple types of annotation in gene-based analysis provides several advantages over analysis methods using annotations of a single type. First, including variants from multiple annotation categories is expected to increase accuracy (e.g., odds that the most significant gene at a locus is causal), since signals that overlap a single annotation type (e.g., eVariants) may be driven by linkage disequilibrium (LD) or pleiotropic regulatory effects (Wainberg, Sinnott-Armstrong, D Knowles, et al. 2017; Wainberg, Sinnott-Armstrong, Mancuso, et al. 2019). Second, it can increase power by increasing the signal-to-noise ratio, and capturing a wider range of possible mechanisms driving genetic associations with complex traits (e.g., AJ Schork et al. 2013; Lu et al. 2016; Kichaev et al. 2019). For example, tests that incorporate both coding variants and eVariants are expected to have high power to detect both protein-altering associations as well as associations driven by effects on gene expression levels. One-dimensional annotation scores derived from multiple annotation data sets can be used to weight variants in gene-based tests (e.g., D Lee et al. 2015; Kelley, Snoek, and Rinn 2016; Rentzsch et al. 2018); however, aggregating variants separately for multiple annotation types and combining the result allows us to explicitly model multiple distinct genes and biological mechanisms underlying associations.

Here, we present a statistical framework and computational tool to integrate heterogeneous functional annotations with GWAS association summary statistics for gene-based analysis. We analyze a diverse set of functional annotation data including multiple tissue-specific eQTL annotation data sets, multiple epigenetic annotation sets mapping regulatory elements to putative target genes, coding variant annotations, and proximity-based annotations. We compare the performance of single-annotation, omnibus, and annotation-agnostic gene-based analysis methods through simulation studies, and by analyzing GWAS summary statistics from the UK Biobank (Bycroft et al. 2018). Our contributions are to 1) expound a general statistical framework for gene-based analysis with heterogeneous functional annotations, which includes several existing single-annotation gene-based association methods as components or special cases; 2) provide a computationally efficient open-source tool for gene-based analysis from summary statistics; and 3) conduct a comprehensive analysis of statistical power and accuracy identifying causal genes across gene-based association methods through extensive simulation studies and analysis of GWAS data for 128 human traits.

## Results

We first outline a statistical framework and open-source tool for gene-based analysis with heterogeneous functional annotations. Next, we describe simulations to evaluate 1) the Type I error rates of gene-based test statistics, 2) statistical power, and 3) specificity to identify causal genes. Finally, we discuss applications to empirical data using GWAS summary statistics from the UK Biobank. We assess 1) the empirical power of gene-based tests by comparing the numbers of significant independent gene-based associations discovered for each UK Biobank trait, and 2) concordance with benchmark gene lists compiled from the ClinVar database (Landrum et al. 2015) and the Human Phenotype Ontology (HPO) (Köhler et al. 2016).

### GAMBIT Framework

GAMBIT (Gene-based Analysis with oMniBus, Integrative Tests) is an open-source tool for calculating and combining annotation-stratified gene-based tests using GWAS summary statistics (single-variant association z-scores). Broadly, GAMBIT’s strategy is to first separately calculate single-annotation gene-based association tests stratified by functional annotation class, and aggregate across classes for each gene to construct omnibus gene-based tests (illustrated in Figure 1). Here and elsewhere, we refer to this omnibus test statistic as the GAMBIT gene-based test. GAMBIT calculates four general forms of gene based test statistics, described briefly in Table 1 and detailed in Materials and Methods. To account for LD between neighboring variants and genes, GAMBIT relies on an LD reference panel from an appropriately matched population (e.g., International HapMap 3 Consortium 2010; 1000 Genomes Project Consortium 2015). GAMBIT is implemented in C++, open source, and freely available.

**Table 1:**
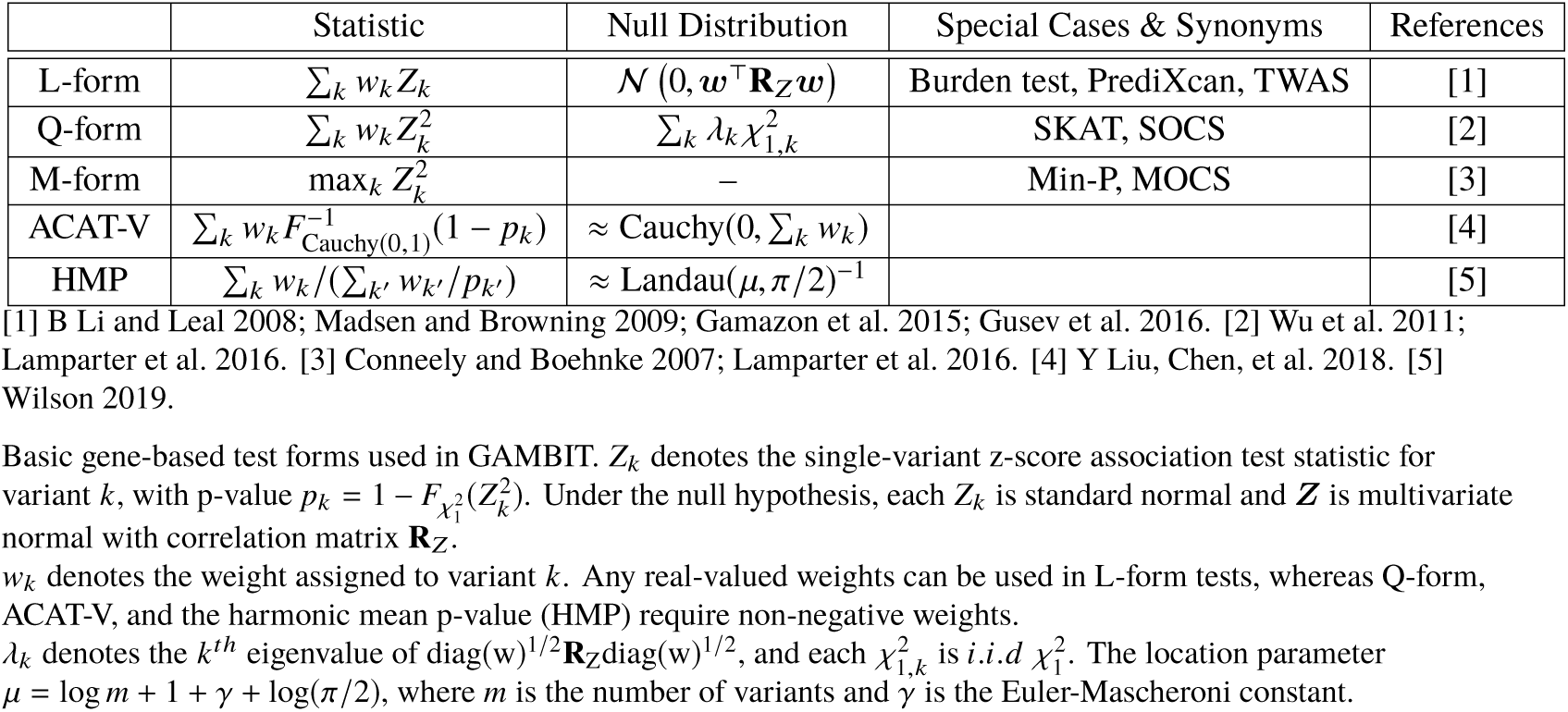
Forms of Gene-Based Test Statistics

|  | Statistic | Null Distribution | Special Cases & Synonyms | References |
| --- | --- | --- | --- | --- |
| L-form | $\sum_k w_k Z_k$ | $\mathcal{N}(0, \mathbf{w}^\top \mathbf{R}_Z \mathbf{w})$ | Burden test, PrediXcan, TWAS | [1] |
| Q-form | $\sum_k w_k Z_k^2$ | $\sum_k \lambda_k \chi_{1,k}^2$ | SKAT, SOCS | [2] |
| M-form | $\max_k Z_k^2$ | — | Min-P, MOCS | [3] |
| ACAT-V | $\sum_k w_k F_{\text{Cauchy}(0,1)}^{-1}(1 - p_k)$ | $\approx \text{Cauchy}(0, \sum_k w_k)$ | | [4] |
| HMP | $\sum_k w_k / (\sum_{k'} w_{k'} / p_{k'})$ | $\approx \text{Landau}(\mu, \pi/2)^{-1}$ | | [5] |

[1] B Li and Leal 2008; Madsen and Browning 2009; Gamazon et al. 2015; Gusev et al. 2016. [2] Wu et al. 2011; Lamparter et al. 2016. [3] Conneely and Boehnke 2007; Lamparter et al. 2016. [4] Y Liu, Chen, et al. 2018. [5] Wilson 2019.
Basic gene-based test forms used in GAMBIT. $Z_k$ denotes the single-variant z-score association test statistic for variant $k$ , with p-value $p_k = 1 - F_{\chi_1^2}(Z_k^2)$ . Under the null hypothesis, each $Z_k$ is standard normal and $\mathbf{Z}$ is multivariate normal with correlation matrix $\mathbf{R}_Z$ .
$w_k$ denotes the weight assigned to variant $k$ . Any real-valued weights can be used in L-form tests, whereas Q-form, ACAT-V, and the harmonic mean p-value (HMP) require non-negative weights.
$\lambda_k$ denotes the $k^{th}$ eigenvalue of $\text{diag}(\mathbf{w})^{1/2} \mathbf{R}_Z \text{diag}(\mathbf{w})^{1/2}$ , and each $\chi_{1,k}^2$ is *i.i.d* $\chi_1^2$ . The location parameter $\mu = \log m + 1 + \gamma + \log(\pi/2)$ , where $m$ is the number of variants and $\gamma$ is the Euler-Mascheroni constant.

**Figure 1:**
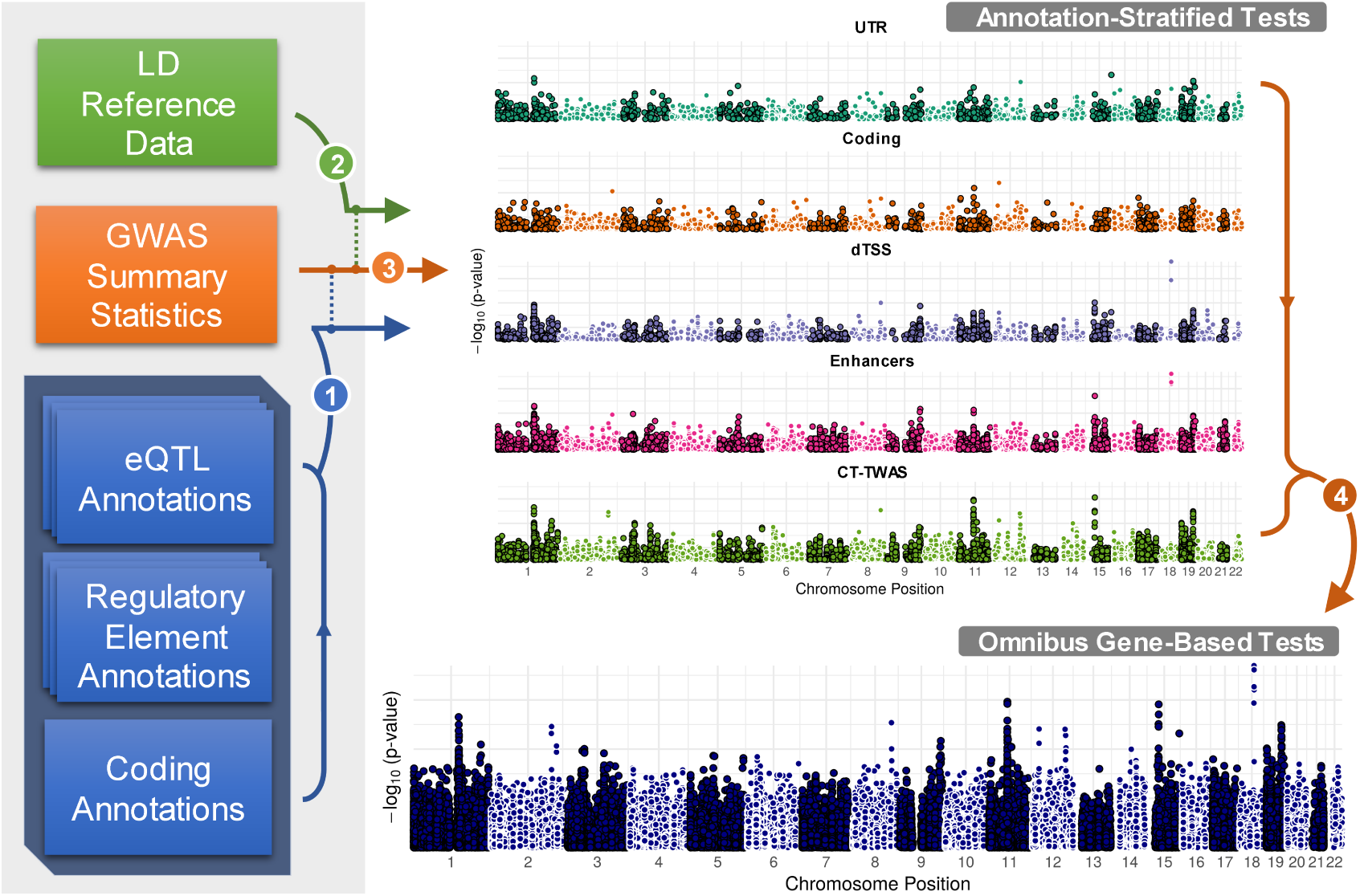
GAMBIT Analysis Framework & Workflow. Broad overview of GAMBIT workflow. (1) GWAS association summary statistics (single-variant z-scores, or effect size estimates and standard errors) are cross-referenced and linked with multiple sets of functional annotations. (2) Annotated GWAS variants are cross-referenced with variants in a haplotype reference panel to estimate LD on-the-fly as needed. (3) GWAS summary statistics, annotations, and LD estimates are used to calculate stratified gene-based test statistics. (4) Stratified gene-based tests are combined for each gene to construct omnibus test statistics. GAMBIT supports multiple single-annotation test methods and multiple omnibus test methods to combine single-annotation tests; detailed statistical methods are provided in Materials and Methods.

### Functional Annotation Data

We considered 5 broad annotation classes in our analysis: 1) proximity-based annotations, 2) coding annotations, 3) UTR regions, 4) enhancer and promoter regions, and 5) eVariants. Each of these annotation classes comprises multiple subclasses; for example, coding annotations include non-synonymous, splice-site, and other variant categories; and eVariants are stratified by tissue. Briefly, we annotated coding and UTR variants using TabAnno (Zhan and DJ Liu 2013) and EPACTS (H Kang 2014); obtained enhancer element and enhancer-target gene weight annotations from RoadmapLinks (Ernst et al. 2011; Kundaje et al. 2015), GeneHancer (Fishilevich et al. 2017), and JEME (Cao et al. 2017); and pre-computed tissue-specific eVariants annotations from PredictDB (Gamazon et al. 2015; A Barbeira et al. 2016) and FUSION/TWAS (Gusev et al. 2016). Enhancer annotations were largely derived from NIH Roadmap Epigenomics and ENCODE project data (Bernstein et al. 2010; ENCODE Project Consortium 2012), as well as from the FANTOM Consortium (Lizio et al. 2015; Marbach et al. 2016; Cao et al. 2017). All eVariant annotations were estimated using the GTEx project v7 data (GTEx Consortium 2015). Figure 2 illustrates a subset of these annotations at the *CELSR2* locus on chromosome 1; detailed descriptions of annotation data and statistical methods used to aggregate test statistics within and across classes are provided in Materials and Methods.

**Figure 2:**
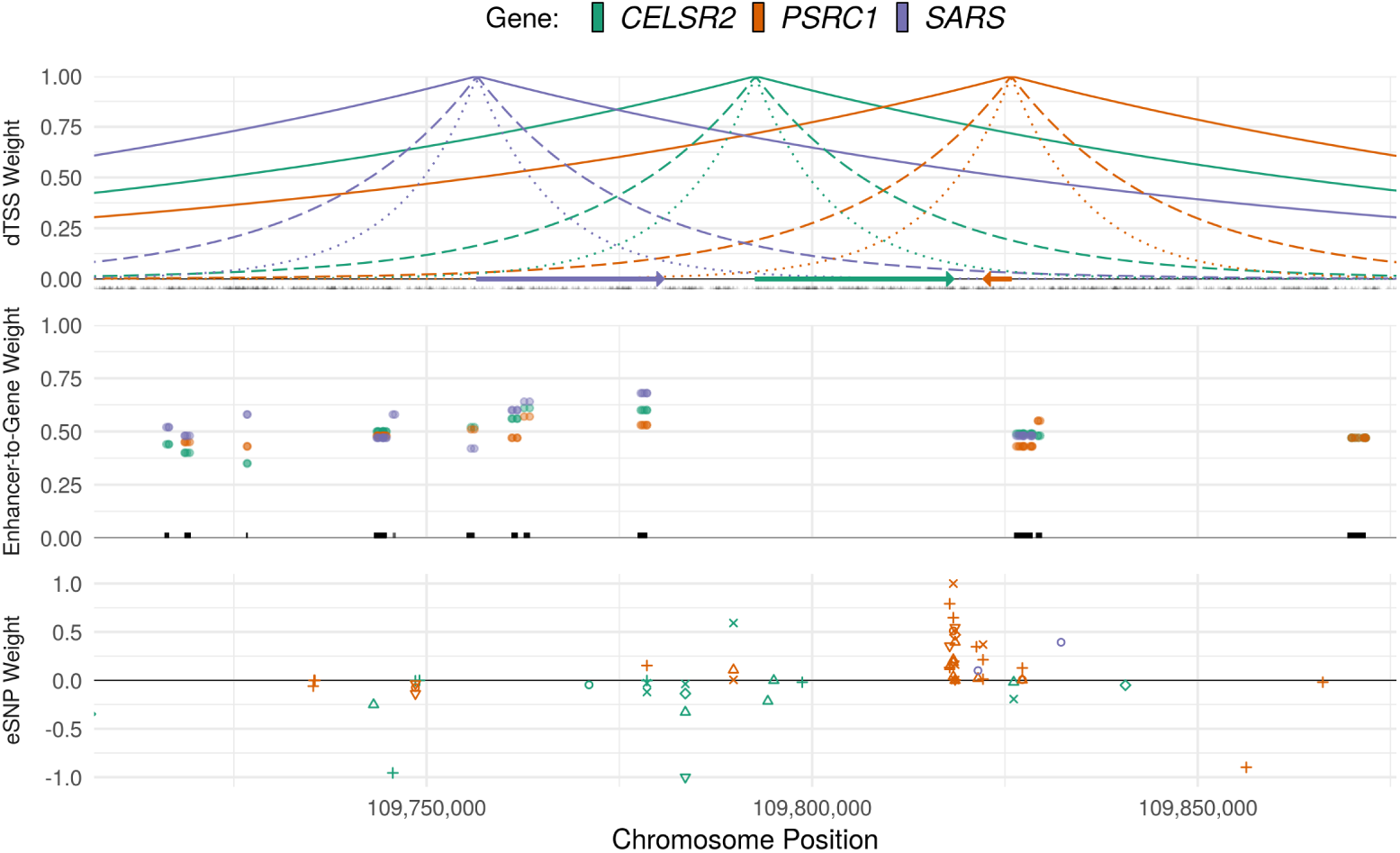
Regulatory Annotation Tracks and Gene Weights. Illustration of primary regulatory annotation tracks used in GAMBIT gene-based analysis framework at the *CELSR2* locus on chromosome 1. Top panel: Distance-to-transcription start site (dTSS) weights, calculated as *w*_*jk*_ (*α*)= exp (− *α* |*d*_*jk*_|), where *d*_*jk*_ is the number of base pairs between variant *j* and the TSS of gene *k*, shown for *α* = 10^−5^ (solid lines), *α* = 5 × 10^−5^ (dashed lines), and *α* = 10^−4^ (dotted lines). Gene bodies are indicated by arrows and variant locations are marked in black at *y* = 0. Middle panel: enhancer-to-target-gene confidence weights. Weights are shown for enhancer variant and target gene, and unique enhancer elements are marked by black lines at *y* = 0. Lower panel: tissue-specific eVariant weights for each gene. eVariant tissues are differentiated by shape.

### GWAS Simulations

We simulated GWAS summary statistics at 2,000 loci using haplotype data from the European subset of the 1000 Genomes Project (1KGP) Phase 3 reference panel (1000 Genomes Project Consortium 2015). Briefly, each locus was defined by first sampling a single causal protein-coding gene, aggregating all genes within 1 Mbp of the causal gene, and finally aggregating all variants assigned to one or more genes based on functional annotations or within ≤ 500kbp of any gene at the locus. For each of the 2,000 loci, we simulated genetic effects under four causal scenarios: coding variants are causal, 2) eVariants are causal, 3) enhancer variants are causal, and 4) UTR variants are causal. For each locus and causal scenario, we varied the proportion of trait variance accounted for by variants at the locus 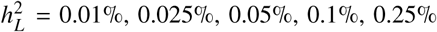 with constant GWAS sample size *n* = 50,000; and for each locus-scenario-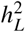 combination, we generated 100 independent simulated replicates. To evaluate p-value calibration and Type I error rates of gene-based tests, we further simulated genome-wide summary statistics for 1,000 traits under the null hypothesis. Detailed simulation procedures are provided in Materials and Methods.

### Simulation Studies: Power and Accuracy Identifying Causal Genes

We compared performance identifying causal genes across 8 gene ranking methods: 1) ranking each gene by distance between its transcription start site (TSS) and the most significant independent single variant at the locus, 2) the Pascal SOCS test -log_10_p-value, which assigns equal weight to all variants within 500kbp of the gene body, 3) the GAMBIT omnibus test -log_10_p-value (labeled as GAMBIT), and 4-8) -log_10_p-values for gene-based tests using each annotation class individually (listed in Table 2 and described in Materials and Methods). As expected, test statistics calculated using the causal annotation class alone were most accurate for identifying the causal gene (e.g., gene-based p-values using coding variants were most accurate when coding variants were causal); however, the GAMBIT omnibus test was nearly as accurate, and had the second-highest performance across simulation settings (Figure 3; Supplementary Figure 1). In practical applications, the causal mechanisms underlying associations are unknown and often heterogeneous across loci; in this case, we expect the GAMBIT omnibus testing strategy to be most accurate.

**Table 2:** Single-Annotation Gene-Based Tests

|  | Test Form | Annotation Subclasses | Annotated Variants |
| --- | --- | --- | --- |
| dTSS | ACAT-V/HMP | dTSS-α value | Variants within 500kbp of TSS |
| CT-TWAS | L-form | eQTL tissue | eVariants across 48 tissues |
| Enhancers | Q-form; ACAT-V/HMP | Enhancer region | All enhancer variants |
| UTR | Q-form; ACAT-V/HMP | 3' and 5' UTR | 3' and 5' UTR variants |
| Coding | Q-form; ACAT-V/HMP | Variant type (e.g., missense, splice site) | Exonic variants |

Summary of variant types, default test methods, and default aggregation procedures for primary annotation classes in GAMBIT. Rationale and further details are provided in Materials and Methods.

**Figure 3:**
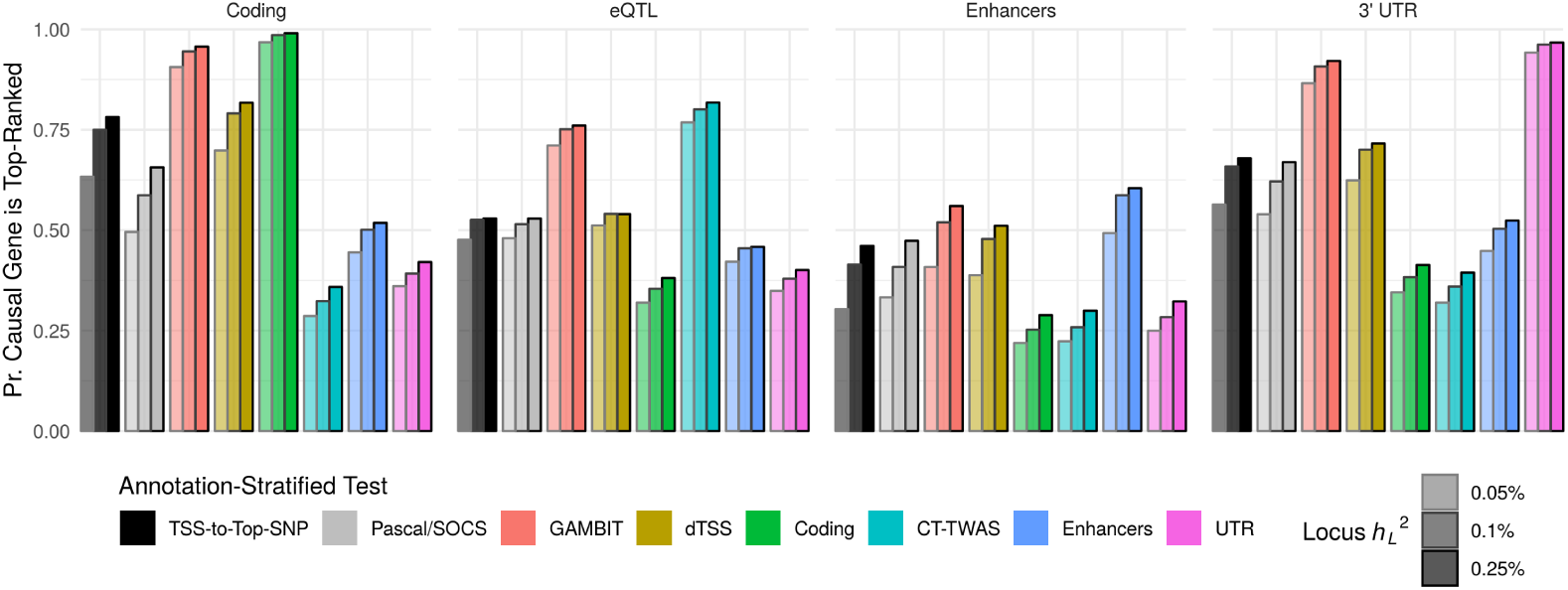
GWAS Simulations: Performance Identifying Causal Gene. Proportion of simulation replicates in which causal gene is top-ranked at at locus (*y*-axis) for each gene-based association or gene ranking method (*x*-axis & bar fill color) stratified by locus heritability 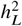 (bar outline color) when either coding, eQTL, enhancer, or UTR variants are causal (plot facet).

We also compared statistical power for each of the gene-based test methods at both causal and non-causal proximal genes at each simulated locus (Figure 4). For proximal genes, association signals are driven by LD and pleiotropic regulatory variants shared with the causal gene; thus, gene-based tests should ideally have high power for causal genes but comparatively low power for proximal genes. Similar to the previous analysis, gene-based tests using the causal annotation class alone had the highest power for causal genes and highest specificity (low power for proximal genes) across simulation settings. The omnibus test generally had the second-highest power for causal genes, and intermediate power for proximal genes. Thus, we expect the omnibus testing approach to be powerful and robust when causal mechanisms are unknown or heterogeneous across loci.

**Figure 4:**
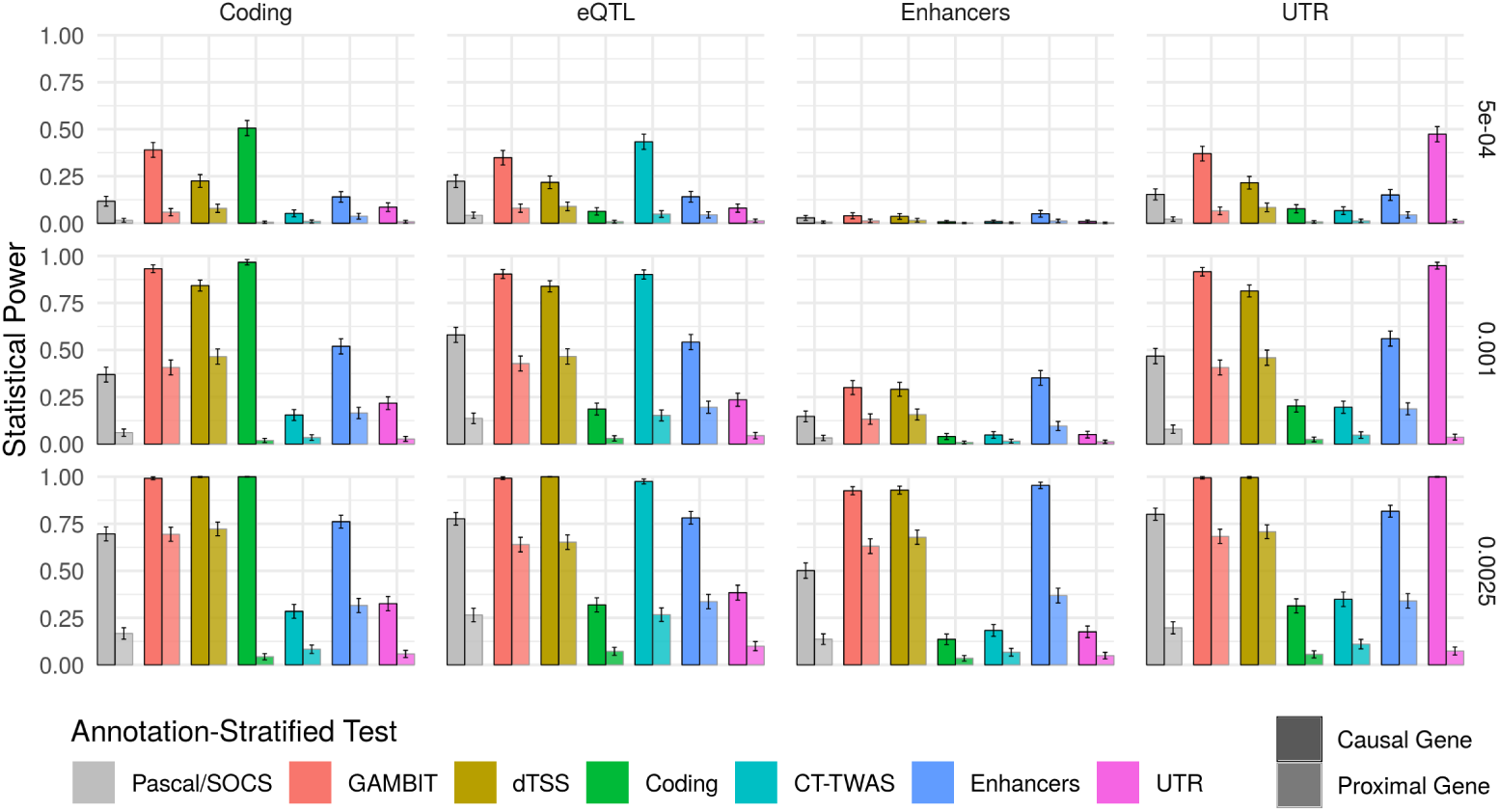
GWAS Simulations: Statistical Power Statistical power. (proportion of simulation replicates in which gene-based *p*-value ≤ 2.5 × 10^−6^ across loci; *y*-axis) for each gene-based testing approach (*x*-axis & color) stratified by locus heritability 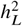 (plot rows) when either coding, eQTL, enhancer, or UTR variants are causal (plot columns). Power is shown separately for causal genes and proximal genes (non-causal genes that are proximal to a causal gene, as defined in Materials and Methods). Ideally, gene-based tests should have high power for causal genes, and relatively lower power for proximal genes. Error bars show 95% confidence intervals for average power across loci.

### Analysis of GWAS Data from the UK Biobank

**Figure 5:**
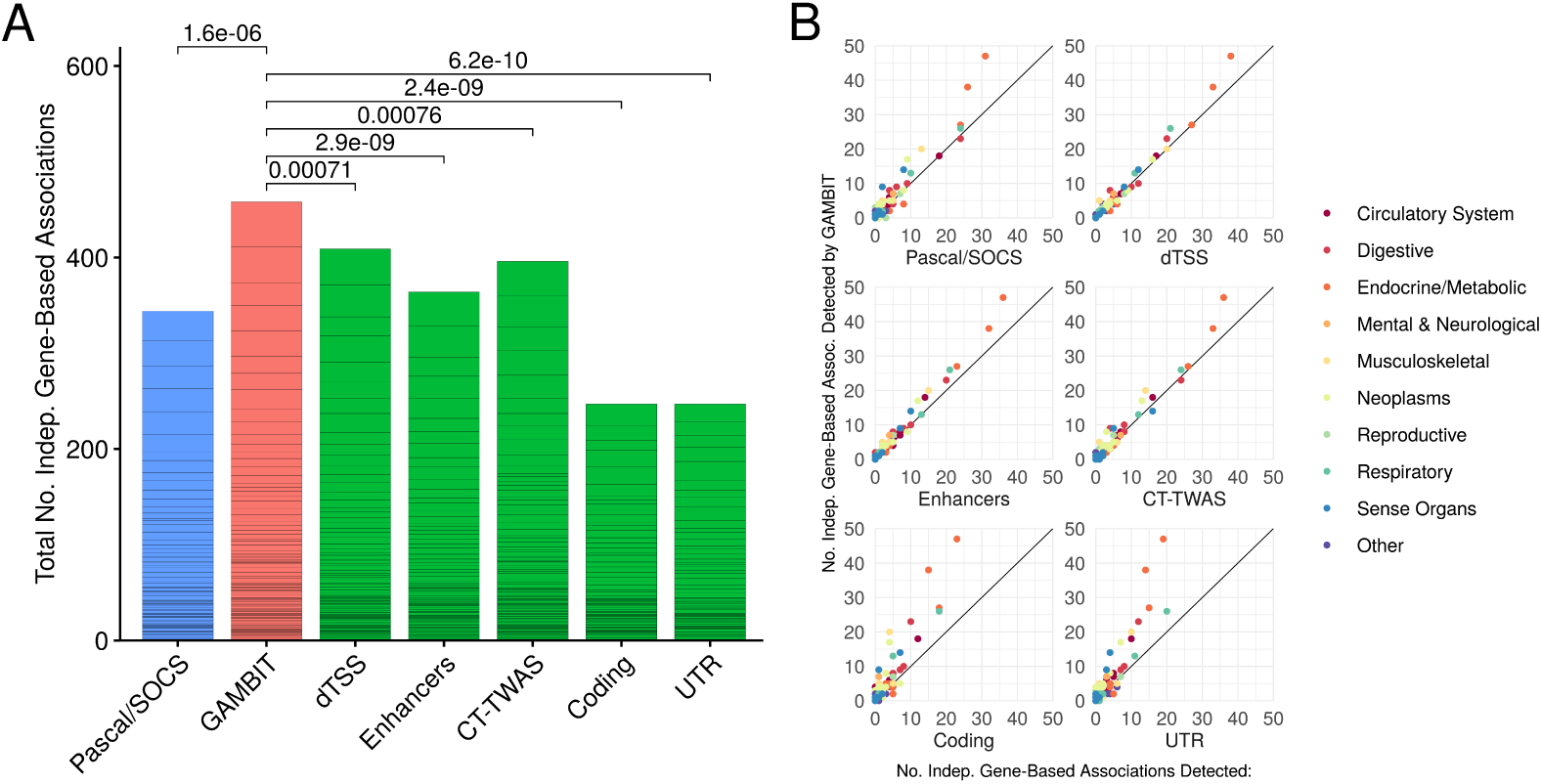
UK Biobank Analysis: Numbers of Significant Independent Associations Detected. Numbers of independent gene-based associations (at Bonferroni-corrected 5% significance level) detected by each method across 128 UK Biobank traits. Panel A: Total number of significant independent associations across traits (delineated by horizontal black lines) for each gene-based test; Wilcoxon signed-rank p-values (top) for paired comparisons between no. associations detected by omnibus test (red) versus Pascal/SOCS (blue) and single-annotation gene-based tests (green). The omnibus test detects significantly more associations than any individual constituent gene-based test or by Pascal/SOCS across UK Biobank traits.

**Figure 6:**
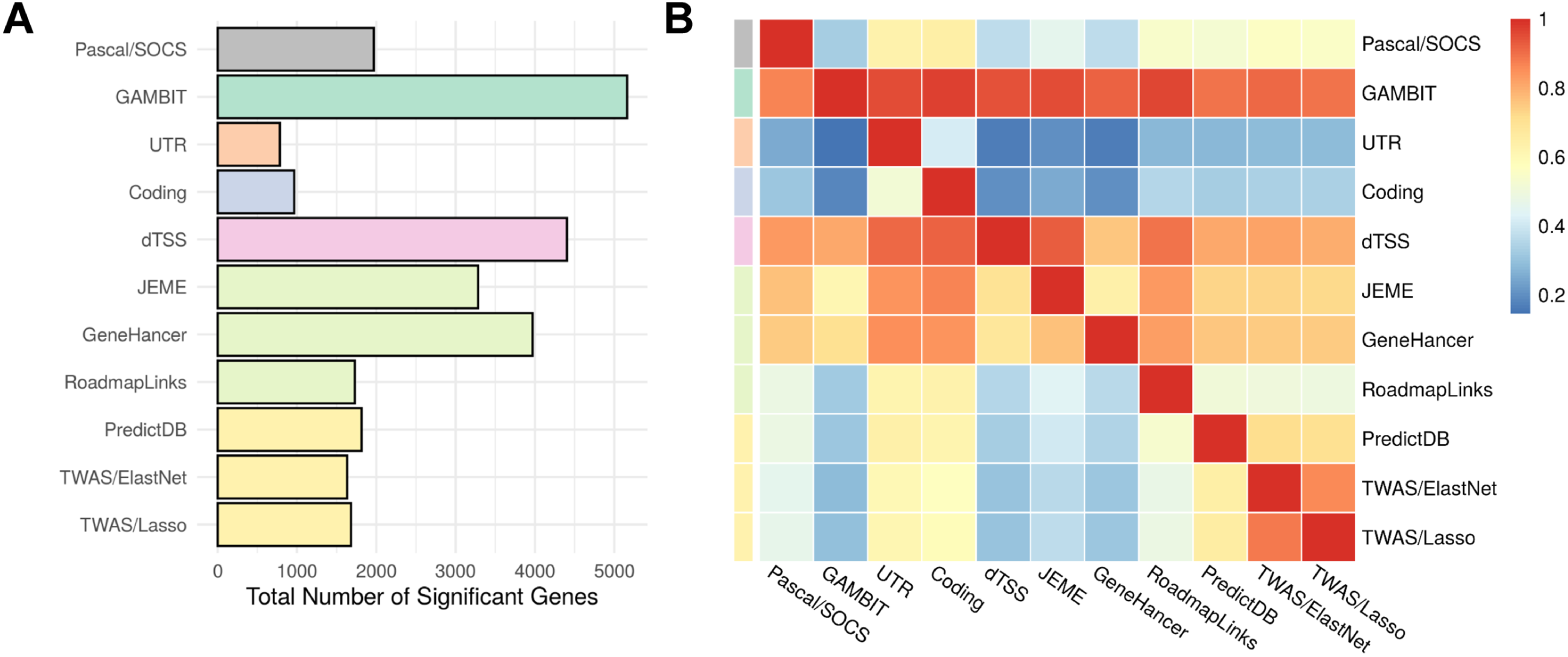
UK Biobank Analysis: Overlap Between Gene-Based Association Methods. Panel A: Total number of significant genes (p-value < 2.5e-6) for each method across all 128 traits. Unlike Figure 5, gene-based associations in Figure 6 are not filtered or LD pruned, and a single significant GWAS variant can produce multiple significant gene-based associations for a given method. Here, a larger number of significant genes does not necessarily imply greater statistical power. Panel B: The *i, j*^*th*^ heatmap element can be interpreted as the conditional probability that gene-based test *i* is significant given that gene-based test *j* is significant, which is estimated as the total number of overlapping significant genes between tests *i* and *j* divided by the total number of significant genes for test *j*.

### Significant Independent Associations Detected for 128 UK Biobank Traits

To compare the power of gene-based tests in empirical data, we evaluated the numbers of significant independent gene-based associations detected for each method across 128 approximately independent GWAS traits in the UK Biobank (selection procedures are described in Materials and Methods). The number of independent associations is calculated for each trait by selecting the most significant gene-based association p-value, masking all gene-based tests that include variants within 1 Mbp of variants for the selected gene, and repeating until all genes with Bonferroni-adjusted p-value ≤ 5% are either selected or masked. This procedure ensures that all selected genes are separated by at least 1 Mbp, and provides a conservative estimate of the number of significant independent signals. Omnibus tests detected significantly more associations than any other gene-based association method considered (Figure 5A), and consistently detected more associations than other methods across a wide range of traits and genetic architectures (Figure 5B).

### Concordance with Benchmark Genes for 25 UK Biobank Traits

We compiled lists of benchmark genes from the ClinVar database (Landrum et al. 2015) and the Human Phenotype Ontology (HPO) (Köhler et al. 2016) for 25 traits in the UK Biobank to compare the gene-based analysis methods identifying causal genes; procedures and selection criteria are detailed in Materials and Methods. Results are shown separately using the union and intersection of ClinVar and HPO benchmark genes; the latter gene set is expected to have higher specificity, albeit fewer genes. Performance identifying benchmark genes was assessed by ranking genes separately within each benchmark locus for each UK Biobank trait, where a benchmark locus is defined as the set of all genes within 1 Mbp of a genome-wide significant single-variant association that also is within 1 Mbp of a benchmark gene. To compare the performance of gene ranking methods, we calculated fraction of loci at which the top-ranked gene coincides with a benchmark gene (Figure 7) and assessed receiver operating characteristic (ROC) and precision-recall curves for each method (Supplementary Figure 2).

**Figure 7:**
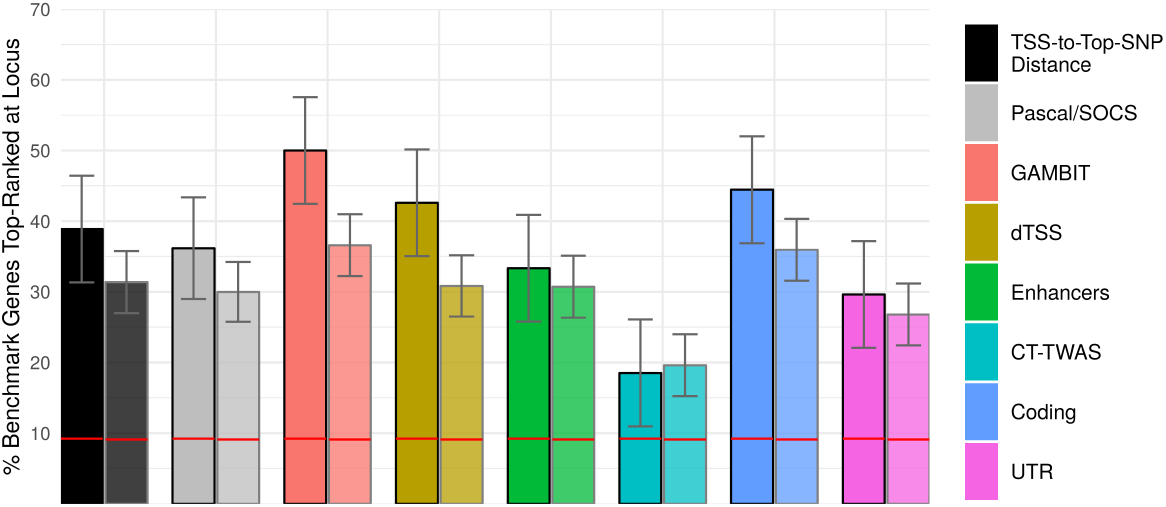
UK Biobank Analysis: Performance Identifying Benchmark Genes. Percentage of loci at which the benchmark gene (identified from HPO and/or ClinVar) is top-ranked for each gene-based association or gene ranking method. For each method, bars on the left (outlined in black) are calculated for benchmark loci present in both HPO and ClinVar (54 loci), and bars on the right (faded outline) are calculated using the union of all HPO and ClinVar loci (153 loci). Horizontal red lines indicate the expected percentage of top-ranked benchmark genes under the null hypothesis that gene rank and benchmark labels are independent. Error bars indicate 95% confidence intervals.

GAMBIT omnibus tests had the highest performance identifying benchmark genes among the gene ranking methods considered, particularly for the stricter gene set, although the difference was not statistically significant relative to most other gene ranking methods (Figure 7). Gene-based tests using coding variants alone had the second-highest performance, which may reflect the enrichment for coding associations within the benchmark gene set (Supplementary Figure 3) caused by benchmark gene selection criteria (described in Materials and Methods). Due to the over-representation of coding associations, Figure 7 may underestimate the impact of incorporating heterogeneous regulatory annotations for associated loci without an established benchmark gene.

Further inspection revealed a number of loci of biological or clinical interest. In the analysis of skin cancer in the UK Biobank, three melanin or melanogenesis-related genes (*TYR, OCA2*, and *MC1R*) and telomerase reverse transcriptase (*TERT*) were top-ranked by GAMBIT, but not top-ranked based on TSS-to-top-SNP distance, while all other benchmark genes for skin cancer were top-ranked by both methods or by neither. At the *TERT* locus, the lead GWAS variant was intronic, whereas the lead variants for *TYR, OCA2*, and *MC1R* were nonsynonymous. Unsurprisingly, the latter three benchmark genes were also top-ranked based on coding variant gene-based p-values; however, only *TERT* was top-ranked based on CT-TWAS.

Similarly, *APOB*, which encodes an apolipoprotein and is associated with autosomal dominant forms of hypercholesterolemia, was top-ranked by GAMBIT but not by TSS-to-top-SNP distance for disorders of lipoid metabolism in the UK Biobank. Despite being >150 Kbp from the intergenic lead GWAS variant, *APOB* was also top-ranked by all single-annotation gene-based tests individually. Conversely, *TSHR*, which encodes a thyroid horomone receptor, was top-ranked based on TSS-to-top-SNP distance but not by GAMBIT for thyrotoxicosis. In this case, the lead GWAS variant was intronic, and CT-TWAS was the only single-annotation gene-based test that ranked *TSHR* as the top gene at its locus. A complete table of results for benchmark genes is provided in Supplementary Materials.

## Discussion

Here, we introduced GAMBIT, a statistical framework and software tool for gene-based analysis with heterogeneous annotations. Our work makes several contributions to the field:

First, we conducted extensive simulation studies to systematically compare gene-based test methods across a range of plausible biological scenarios, and demonstrated pitfalls of test methods that use only a single annotation class. When causal mechanisms are misspecified (i.e., causal variants do not overlap annotated variants used in gene-based analysis), standard gene-based tests have limited power, and can be confounded by LD and pleiotropic regulatory variants that affect multiple genes. This may lead researchers to misidentify the genes and biological mechanisms that contribute to disease risk. Finemapping, co-localization, and conditional analysis can be applied to refine association signals and mitigate spurious inferences following gene-based analysis (e.g., Giambartolomei et al. 2014; Z Zhu et al. 2016; Y Lee et al. 2018; Mahajan et al. 2018). By contrast, our omnibus testing strategy helps to ameliorate spurious inferences within the context of gene-based testing directly, and also has high power to detect associations across a range of causal mechanisms underlying genetic associations.

Second, we analyzed 128 traits from the UK Biobank to evaluate performance in empirical data across a range of complex traits and genetic architectures, and confirmed that incorporating annotations of many types and across many tissues increases power relative to standard methods. While our analysis of concordance with gold-standard causal genes was limited by the relatively small numbers of benchmark genes identified for UK Biobank traits and the inherent difficulty establishing causal genes underlying regulatory associations, we found suggestive evidence that incorporating diverse annotation types in gene-based analysis can improve performance identifying causal genes relative to standard approaches (e.g., ranking genes by distance to the most significant single variant) and gene based tests using a single annotation type.

Finally, we provide a unifying framework and easy-to-use software tool to incorporate heterogeneous functional annotations in gene-based analysis. From its inception, gene-based analysis was built on the premise that aggregating functional variants at the gene level can increase statistical power and help identify causal genes in GWAS (Neale and Sham 2004). Early gene based test methods were developed primarily for rare genic variants (e.g., B Li and Leal 2008; Madsen and Browning 2009), and early gene-based association analyses often used only deleterious coding variants (e.g., Purcell et al. 2014; Majithia et al. 2014). However, functional genomics studies have shown that most functional variation is non-coding (ENCODE Project Consortium 2012), and most variant associations discovered through GWAS to date occur in non-coding regions (Welter et al. 2013; MacArthur et al. 2016), highlighting the importance of regulatory annotations for gene-based association analysis. The first gene-based tests developed explicitly for regulatory variation were TWAS and PrediXcan, which aggregate eVariants to construct proxy variables for tissue-specific gene expression levels using predictive weights estimated from external eQTL mapping data (Gusev et al. 2016; A Barbeira et al. 2016). However, functional and regulatory genomics projects have introduced a wealth of annotations with potential utility for gene-based analysis (e.g., Lizio et al. 2015; Cao et al. 2017; Fishilevich et al. 2017; Stranger et al. 2017).

Our omnibus testing strategy is expected to perform best when variants from a single annotation class (e.g., coding variants) are causal at a given locus. When multiple independent signals from different annotation classes exist at a single gene locus, this testing strategy is expected to have lower power than one that explicitly accounts for multiple possible signal sources (e.g., via Barnett, Mukherjee, and X Lin 2017). While we did not explore this possibility in our simulations, it is an interesting question which we defer to future work.

The utility of incoporating annotations in gene-based analysis depends crucially on the accuracy and comprehensiveness of the underlying annotation data sets. While we considered the case that causal variants may be misspecified, our simulations assumed that the confidence weights assigned to regulatory elements are well-calibrated, and that causal eVariants are annotated. Violations of these assumptions will reduce both power and accuracy in gene-based analysis, and may in part account for differences between our results with empirical versus simulated data. Current transcriptomic and epigenomic studies are generally limited to a subset of human tissues and cell-types, and are derived from data sets of limited sample size (e.g., Stranger et al. 2017; ENCODE Project Consortium 2012). Thus, we expect current transcriptomic and epigenomic annotations to be incomplete and imprecise. Looking forward, larger and more comprehensive studies will enable more comprehensive and accurate annotations, increasing the utility of annotation-informed association analysis methods.

In summary, our work builds upon and generalizes previous gene-based association methods, providing a flexible framework for gene-based analysis with heterogeneous annotations that can be readily adapted when new annotation resources are developed and released.

## Materials and Methods

We describe 1) methods to aggregate variants for gene-based analysis, 2) procedures to combine multiple gene-based tests, 3) functional genomic and annotation data resources, 4) procedures to simulate GWAS data using real genotype and functional annotation data, and 5) GWAS data from the UK Biobank to which we applied our methods.

### Multiple-Variant Association Test Statistics

Here, we review statistical methods to aggregate multiple variants for gene-based, region-based, or pathway association analysis. For convenience, we assume a quantitative trait and ignore the presence of covariates; however, our results can easily be adapted to other settings.

#### Linear-Form Gene-Based Tests (L-form)

The oldest and most widely used gene-based tests are linear combinations of genotypes across variants (B Li and Leal 2008; Madsen and Browning 2009; S Lee, Wu, and X Lin 2012), here referred to as L-form tests. Formally, we define the L-form test as *T*_*L*_ = (***w***^T^**R**_*Z*_ ***w***)^−1/2^***w***^T^***Z***, where ***w*** is a vector of single-variant weights, ***Z*** is a vector of single-variant association statistics (where each *Z*_*j*_ follows the standard normal distribution under the null hypothesis), and **R**_*Z*_ is the correlation matrix of z-scores. Under the null hypothesis of no association, *T*_*L*_ follows the standard normal distribution. The L-form test statistic *T*_*L*_ can be computed from GWAS summary statistics (single-variant z-scores, or effect sizes and standard errors) and covariance estimates, and can be written either as linear combinations of single-variant association statistics or as linear combinations of genotypes (ZZ Tang and DY Lin 2013; DJ Liu et al. 2014).

Examples of L-form tests include burden tests, which calculate burden scores as a weighted sum of rare, putatively deleterious mutations (B Li and Leal 2008; S Lee, Wu, and X Lin 2012); and TWAS/PrediXcan tests (Gamazon et al. 2015; Gusev et al. 2016; A Barbeira et al. 2016), which aggregate eQTL variants using predictive weights estimated from external eQTL mapping data, e.g. from the GTEx project (GTEx Consortium 2015). These can be viewed as tests of association between GWAS trait and an explicit proxy variable constructed as a linear combination of genotypes. Importantly, L-form tests rely on prior knowledge regarding the directions of effect across variants (Wu et al. 2011; S Lee, Wu, and X Lin 2012). For example, the signed weights used in burden tests often reflect the hypothesis that rare deleterious alleles increase risk for disease, and the predictive weights used in TWAS/PrediXcan reflect the hypothesis that gene expression mediates the associations between genotypes and complex trait.

#### Quadratic-Form Gene-Based Tests (Q-form)

Variance component tests and quadratic forms of single-variant association statistics comprise another widely used class of gene-based association methods, here referred to as Q-form (quadratic) tests. Q-form tests include VEGAS (or SOCS), defined as the sum of squared single-variant z-scores (JZ Liu et al. 2010; Lamparter et al. 2016); and SKAT, a weighted quadratic form of single-variant association statistics (Wu et al. 2011). Formally, the Q-form test statistic is defined *T*_*Q*_ = ***Z***^T^diag(***w***)***Z***, where diag(***w***) is a diagonal weight matrix and ***Z*** is a vector of single-variant association z-scores; under the null hypothesis of no association, *T*_*Q*_ follows a mixture chi-squared distribution with mixture proportions equal to the eigenvalues of diag(***w***)^1/2^**R**_*Z*_ diag(***w***)^1/2^, where **R**_*Z*_ is the correlation matrix of z-scores. In contrast to L-form tests, Q-form tests aggregate single-variant association statistics without prior knowledge or assumptions pertaining to the directions of effects across variants (Wu et al. 2011; S Lee, Wu, and X Lin 2012). While less tractable than L-form, analytical p-values for Q-form tests can be calculated using a variety of techniques to approximate the tail probabilities of multivariate normal quadratic forms (e.g., Davies 1980; H Liu, Y Tang, and HH Zhang 2009), which are far more efficient than permutation procedures or Monte Carlo methods (Mishra and Macgregor 2015; Lamparter et al. 2016). Q-form tests are most appropriate when a sizable proportion of variants are hypothesized to have non-zero effects of unknown and inconsistent direction (S Lee, Wu, and X Lin 2012).

#### Maximum Chi-Squared Statistic as a Gene-Based Test (M-form)

Perhaps the simplest gene-based test is the maximum chi-squared statistic across variants (or equivalently, the minimum p-value), here referred to as M-form tests. Analytical p-values for M-form tests can be calculated by directly integrating the multivariate normal density of z-scores within the hypercube given by ***x*** ∈ ℝ^*m*^ : max_*k*_ | *x*_*k*_ | ≤ max_*j*_ | *Z*_*j*_ | where *m* is the number of variants, or approximated by adjusting the minimum p-value across variants by the effective number of tests (Conneely and Boehnke 2007; Lamparter et al. 2016). M-form tests are most appropriate when only one or a small fraction of variants are hypothesized to have non-trivial effects.

#### Aggregated Cauchy Association Test (ACAT)

The aggregated Cauchy association test (ACAT), a recently proposed method to combine multiple dependent p-values, can be used to construct gene-based tests by transforming single-variant association p-values using the Cauchy quantile and cumulative distribution functions, and computing a p-value)

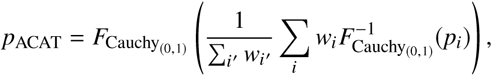

where *p*_*i*_ and *w*_*i*_ are the p-value and weight for the *i*^*th*^ variant and 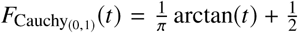 is the CDF of the standard Cauchy distribution (Y Liu, Chen, et al. 2018; Y Liu and Xie 2018). ACAT is expected to perform well when only a small fraction of variants are causal (Y Liu, Chen, et al. 2018). Importantly, ACAT does not require LD computation, and can thus be calculated in *O*(*m*) time where *m* is the number of variants.

#### Harmonic Mean P-value (HMP)

Another recently proposed method to combine multiple dependent p-values, the Harmonic Mean P-value (HMP; Wilson 2019), can similarly be used to construct gene-based tests by weighting p-values from single-variant association tests. The unadjusted HMP p-value is defined

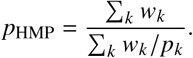

While this statistic can be anti-conservative when directly interpreted as a p-value, Wilson (2019) showed that 1/*p*_HMP_ follows a Landau distribution (with scale and location parameters given in Table 1), which can be used to compute an asymptotically exact HMP p-value. The Landau density function is

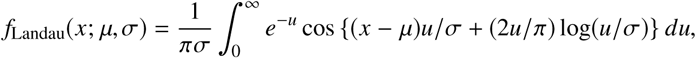

which can be computed numerically with high precision using asymptotic expansions (Kölbig and Schorr 1983). To improve p-value calibration, we implemented the asymptotically exact HMP in the GAMBIT software tool. Unlike L-form and Q-form tests, p-values from M-form, ACAT, and HMP tests are greater than or equal to min_*i*_ *p*_*i*_. However, these methods can still increase power relative to single-variant analysis by reducing the burden of multiple testing and assigning higher weight to functional variants.

#### Generalizations and Extensions

The simple forms of gene-based tests described above can be related and combined through a variety generalizations and extensions. Q-form and M-form can both be viewed as special cases of a statistic (Σ_*j*_ *w*_*j*_ | *Z*_*j*_ |^*p*^)^1/*p*^, which is equivalent to Q-form when *p* = 2 and to M-form when *p* → ∞; this generalization has been used, for example, in the aSPU gene-based test (Kwak and Pan 2015). Similarly, Q-form and L-form can both be viewed as special cases of a statistic 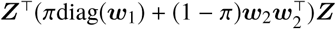, which is equivalent to Q-form when π = 1 and L-form when π = 0; this generalization has been used, for example, in the SKAT-O gene-based test (S Lee, Wu, and X Lin 2012). Finally, ACAT and HMP can be used to combine p-values across multiple gene-based test forms (Y Liu, Chen, et al. 2018; Wilson 2019).

### Integrating Functional Annotations in Gene-Based Tests

#### dTSS Weights

One of the most common heuristics to infer putative causal genes at GWAS loci in the absence of functional annotation is to rank genes by distance between their transcription start site (TSS) and the peak GWAS variant. This strategy is appealing given the strong enrichment of regulatory variants near TSS.

To incorporate distance-to-TSS (dTSS) and capture association signals at regulatory variants that are not well-annotated in gene-based analysis, we define the dTSS weights for gene *k* as 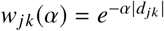, where *d*_*jk*_ is the genomic distance (number of base pairs) between variant *j* and the TSS for the gene of interest. Larger values of the parameter *α* confer more weight to variants nearer the TSS. In practice, we only include variants within a specified window (e.g., 500kbp) of the TSS of the corresponding gene. While dTSS weights can be used in any weighted gene-based test (e.g., Q-form tests), ACAT and HMP are particularly well-suited due to their linear computational complexity, as dTSS-weighted tests often involve thousands of variants per gene.

The optimal *α* value is expected to vary across loci, and likely depends on local gene density and other factors. However, ACAT and HMP can be applied again to calculate an overall p-value that combines dTSS-weighted gene-based test p-values *p*_*k*_ (*α*_*i*_) across multiple values *α*_1_, *α*_2_, …(Y Liu, Chen, et al. 2018; Y Liu and Xie 2018). By default, GAMBIT calculates overall dTSS-weighted test statistics by aggregating across *α* values 10^−4^, 5 × 10^−5^, 10^−5^, 5 × 10^−6^.

#### Regulatory Element-Target Gene Weights

To capture association signals across regulatory elements that have been assigned to one or more target gene, we weight variants in regulatory elements by element-to-target-gene confidence scores, and aggregate variants for each gene using either ACAT, HMP, or Q-form gene-based test statistics. For example, we define the regulatory-element weighted Q-form test statistic as

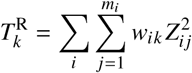

where *m*_*i*_ is the number of variants in the *i*^*th*^ regulatory element, *w*_*ik*_ is the confidence weight between element *i* and gene *k*, and *Z*_*i* *j*_ is the *j*^*th*^ variant in the *i*^*th*^regulatory element.

#### eQTL Weights

Given a vector of weights ***b***_*kt*_ to predict expression levels for gene *k* in a given tissue or cell type *t* as a linear combination of normalized genotypes, the z-score TWAS test of association between predicted expression level and GWAS trait is

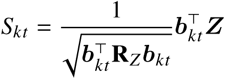

where ***Z*** is the vector of single-variant GWAS z-scores and **R**_*Z*_ is the correlation matrix of z-scores.

To aggregate test statistics across multiple tissues or cell-types, which we refer to as Cross-Tissue TWAS (CT-TWAS), we considered three approaches:

1. Q-form Cross-tissue Test (CT-Q): Calculating the sum of squared tissue-specific test statistics, 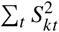, which has a mixture chi-squared distribution under the null hypothesis of no association,
2. M-form Cross-tissue Test (CT-M): Calculating an analytic p-value for the maximum absolute test statistic max_*t*_ |*S*_*kt*_ | using the multivariate normal joint density of tissue- or cell-type-specific test statistics *S*_*k*1_, *S*_*k*2_, … under the null hypothesis of no association, and
3. ACAT or HMP Cross-tissue Test (CT-A or CT-H): Combining tissue- or cell-type-specific p-values *p*_*kt*_ = 2Φ(||*S*_*kt*_ |) using the ACAT method or HMP respectively.

CT-Q and CT-.JM require the cross-tissue correlation matrix **R**_*S*_ with elements 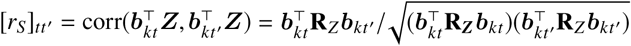, which can be computed in *O*(*m*^2^*n* + *mn*^2^) time where *m* is the number of tissues or cell-types and *n* is the number of eVariants. By contrast, CT-A and CT-H p-values can be computed in *O*(*m*) time, since ACAT and HMP do not require the correlation matrix to be computed. By default, GAMBIT implements CT-A; in our analysis of UK Biobank data, CT-M, CT-H, and CT-A generally perform similarly, while CT-Q tends to detect fewer significant associations.

#### Combining Single-Annotation Test Statistics

In early versions of GAMBIT, we combined gene-based p-values across annotation classes for each gene using standard family-wise error rate (FWER) and false detection rate (FDR) controlling procedures. However, two recently proposed methods, the Aggregated Cauchy Association Test (ACAT; Y Liu and Xie 2018) and Harmonic Mean P-value (HMP; Wilson 2019), provide more powerful approaches to combine multiple dependent p-values, and we have therefore implemented both of these methods in GAMBIT. We calculated ACAT p-values (defined above) using standard formula for the Cauchy quantile function and CDF. To calculate asymptotically exact HMP p-values, we adapted a C++ routine from the ROOT System (Brun and Rademakers 1997) for computing the Landau distribution CDF following the derivations of Wilson (2019) for the asymptotic distribution of the HMP.

### Functional Annotation Data Sources

#### Promoter and Enhancer-Target Annotation Data

To identify regulatory genetic elements and their putative target genes, we used pre-computed annotation data sets from three existing methods: Joint Effects of Multiple Enhancers (JEME) (Cao et al. 2017), GeneHancer (Fishilevich et al. 2017), and RoadmapLinks (Y Liu, Sarkar, et al. 2017; Ernst et al. 2011; Kundaje et al. 2015). GeneHancer provides a global confidence score between each enhancer element and one or more putative target genes, while JEME and RoadmapLinks provide tissue- or cell-type-specific enhancer-target confidence scores. For the latter two data sets, we calculated overall enhancer-target confidence scores across tissues and cell types as the soft maximum (*LogSumExp* function) of tissue- or cell-type-specific scores for each enhancer-target pair. Descriptive statistics for each enhancer annotation dataset are provided in Supplementary Table 2.

#### Tissue-Specific eVariant Annotations and Predictive Weights

To incorporate eVariants in gene-based analysis, we used pre-computed tissue-specific predictive weights for eGene expression estimated using GTEx v7 (Stranger et al. 2017) from TWAS/FUSION (including elastic net and LASSO models) (Gusev et al. 2016) and PredictDB (Gamazon et al. 2015; AN Barbeira et al. 2018). We generated a GAMBIT eWeight annotation files incorporating all available tissues and cell types for each data resource and predictive model. Descriptive statistics for each eVariant weight dataset are provided in Supplementary Table 1.

#### Coding Variant and Gene Annotations

We annotated coding variants, TSS locations, and UTR variants using TabAnno 419 (Zhan and DJ Liu 2013) and EPACTS (H Kang 2014) based on GENCODE v14 (Harrow et al. 2012).

### Simulation Procedures

Here, we describe procedures to simulate GWAS summary statistics using real genotype data or LD estimates. We begin by defining summary statistics and deriving their distribution. We next outline procedures to simulate GWAS summary statistics under the desired distribution. Finally, we describe procedures to simulate configurations of causal genes, causal variants, and effect sizes using real functional genomic annotation data.

#### Simulating GWAS Summary Statistics

We simulated GWAS traits under the model

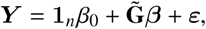

where ***Y*** ∈ ℝ^*n*^ is a quantitative trait for a GWAS sample of size *n*, **1**_*n*_ is the *n* × 1 vector of 1’s and *β*_0_ ∈ ℝ is the trait intercept, 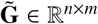 is the centered and scaled genotype matrix where each column has mean 0 and variance 1, ***β*** ∈ ℝ^*m*×1^ is a vector of causal genetic effects, and ***ε*** ∈ ℝ^*n*^ is an *i.i.d.* trait residual with 𝔼(*ε*_*i*_) = 0 and 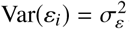.

The total trait variance is

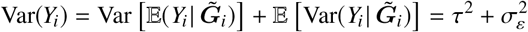

where the genetic variance 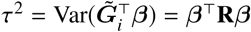, and 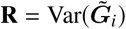 is the genotype correlation (LD) matrix. Here, we treat the genetic effects ***β*** as a constant vector rather than a random variable, and write 𝔼 (·) rather than 𝔼 (·|***β***) to simplify notation. In addition, we fix *β*_0_ = 0 and 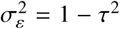 here and elsewhere (without loss of generality). By fixing the residual variance 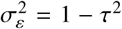, we can interpret *τ*^2^ as the trait heritability, i.e. the proportion of trait variance due to genetic effects.

Single-variant GWAS association analysis aims to detect marginal associations between trait and genotypes at individual variants rather than multiple variants jointly. The marginal effect of variant *k* is

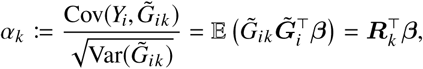

where ***R***_*k*_ is *k*^*th*^ row (or column) of the LD matrix **R**. We note that *α*_*k*_ quantifies a statistical association (marginal covariance) between variant *k* genotypes and trait, which is a function of both *β*_*k*_ and causal effects *β*_*k*′_ for variants *k*′in LD with *k*. The single-variant association test statistic corresponding to the null hypotheses *H*_0_ : *α*_*k*_ = 0 is

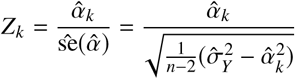

where 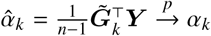 and 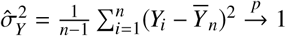.

We can write the vector of single-variant association statistics for variants *k* = 1, 2, …, *m* as

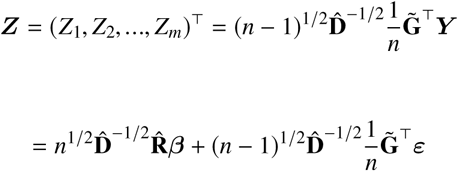

where 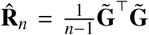 is the sample LD matrix, and 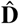 is an *m* × *m* diagonal matrix with 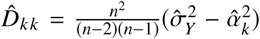. Note that 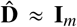 if the proportion of trait variance accounted for by each individual variant is small (e.g., < 1%). If trait residuals ***ε*** are i.i.d. normal with mean 0 and variance 1, then

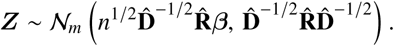

We simulated GWAS summary statistics by calculating 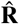 from the European subset of the 1000 Genomes Project panel, and replacing 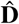 by its limiting value **D** with elements 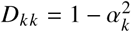. Because 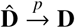, Slutsky’s theorem implies that test statistics calculated using **D** are asymptotically equivalent to those using 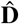, which is acceptable when the GWAS sample size *n* is large.

#### Simulating Genetic Effects at Causal Loci

We used empirical functional annotation data to simulate causal genetic effects ***β*** under realistic genetic architectures. For each simulated causal locus, we selected a causal gene by sampling a single CCDS protein-coding gene, and defined proximal genes as any gene with TSS within 1 Mbp of the causal gene TSS. We then simulated single-variant GWAS summary statistics for all variants associated with any causal and proximal genes by proximity (≤ 1 Mbp) or functional annotations (e.g., eQTL variants).

We simulated causal genetic effects under 5 scenarios: 0) no association (null model), 1) coding association, 2) enhancer association, 3) eGene association, and 4) UTR association. For coding and UTR associations, we first selected the number of causal variants 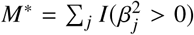 from a Poisson distribution with rate parameter *λ* = *M*/4 truncated to 1 ≤ *M*\* ≤ *M*, where *M* is the total number of coding (or UTR) variants for the causal gene, and randomly selected *M*\* causal variants from the total set of *M* coding (or UTR) variants for the causal gene. This procedure results in ∼25% of all coding (or UTR variants) having non-zero causal effects, while ensuring that at least one variant is causal. For enhancer associations, we similarly simulated the number of causal enhancers *M*\* from a Poisson distribution with rate parameter *λ* = *M*/4, where *M* is the number of enhancers mapped to the causal gene, and selected causal enhancers using a categorical distribution with probability weights derived from confidence scores between enhancer elements and the causal gene. For eGene associations, we selected a single causal tissue at random, and simulated causal effect sizes proportional to precomputed eVariant weights for the causal gene and tissue. Because eVariant weights are noisy in practice, we used simulated eVariant weights 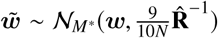 in place of the original eVariant weight vector ***w*** in TWAS gene-based tests, where *N* is the GTEx v7 sample size for the causal tissue.

### The UK Biobank Resource

We used GWAS summary statistics (single-variant association effect size estimates, standard errors, and p-values) for a set of 1,403 traits in the UK Biobank (Bycroft et al. 2018) cohort calculated using SAIGE (Zhou et al. 2018). Genotype data were imputed using the Haplotype Reference Consortium panel (McCarthy et al. 2016), and filtered to include only variants with imputed MAC > 20 in the UK Biobank. We selected a subset of 189 traits for primary analysis by including only traits with effective sample size ≥ 5, 000, and ≥ 1 single-variant association p-value ≤2.5e-8. For our analysis of empirical power, we selected a subset of 128/189 traits by iteratively pruning pairs of correlated traits. Beginning with the most highly correlated pair of traits, we retained the trait with the larger number of significant independent single-variant associations (in the case of ties, we selected the trait with the most detailed description), and repeated this procedure until the maximum pairwise correlation-squared between traits was ≤0.10. For our analysis of concordance with benchmark genes, we first selected a subset of 47 traits including only traits with ≥ 1 single-variant association p-value < 5e-10, excluding benign neoplasms, and including at most a single trait within each trait category. We identified ≥ 1 relevant benchmark genes for 25 of the original 47 traits.

### Selection of Benchmark Genes

To identify benchmark genes for each of the traits selected from the UK Biobank, used the ClinVar (Landrum et al. 2015) and Human Phenotype Ontology (HPO) databases (Köhler et al. 2016). The HPO database explicitly links genes to traits, while the ClinVar database links traits to variants. To identify benchmark genes from ClinVar, we extracted protein-altering variants (frameshift, missense, nonsense, splice site, or stop-loss variants), and excluded variants with unknown or ambiguous molecular consequence (e.g., intergenic and intronic variants). Despite including only ClinVar genes with coding associations, we expect to capture some genes for which both rare coding variants and common regulatory variants contribute to disease risk. For each UK Biobank trait, we extracted all protein-altering ClinVar variants +/-1 Mbp of a genome-wide significant UK Biobank variant, and manually selected ClinVar traits equivalent or closely related to the corresponding UK Biobank trait. We then annotated genes associated with one or more relevant ClinVar trait as a ClinVar benchmark gene. We identified benchmark genes from the HPO database by manually matching keywords between UK Biobank and HPO traits. A complete list of HPO/ClinVar traits and benchmark genes for each UK Biobank trait is provided in Supplementary Materials.

## Data Access

GAMBIT Software: https://github.com/corbinq/GAMBIT

UK Biobank SAIGE Summary Statistics: ftp://share.sph.umich.edu/UKBB_SAIGE_HRC/

### eQTL Annotation Data Sources

PredictDB: http://predictdb.org/

TWAS/FUSION: http://gusevlab.org/projects/fusion/#reference-functional-data

### Enhancer Annotation Data Sources

RoadmapLinks: www.biolchem.ucla.edu/labs/ernst/roadmaplinking

JEME: http://yiplab.cse.cuhk.edu.hk/jeme/

GeneHancer: https://www.genecards.org/

## Acknowledgments

We acknowledge all participants and researchers of the 1000 Genomes Project and UK Biobank study. We thank Xihong Lin for providing valuable comments and suggestions on the manuscript. We thank Sarah Gagliano and Jonas Billie Nielson for suggestions regarding annotations and assistance with UK Biobank data, and Wei Zhou for generating SAIGE single-variant association summary statistics for UK Biobank traits. This research has been conducted using the UK Biobank Resource under Application Number 24460.

## Contributions

CQ and HMK designed the analysis framework. CQ and HMK designed the simulation studies and data analysis. CQ developed the GAMBIT software with assistance from XW. CQ conducted the simulation studies and gene-based analysis of UK Biobank data. CQ generated the figures. CQ and HMK wrote the manuscript. GA suggested the problem. All authors commented on and edited the manuscript. HMK and MB supervised the project.

## Supplementary Tables and Figures

**Supplementary Table 1:** Descriptive Statistics for eQTL Annotation Data Sets

|  | TWAS/FUSION |  | PredictDB |
| --- | --- | --- | --- |
|  | Elastic Net | LASSO | Elastic Net |
| No. Tissues | 48 | 48 | 48 |
| Total No. eVariants | 766,885 | 451,186 | 1,437,864 |
| Total No. eGenes | 25,418 | 25,634 | 25,844 |
| Mean No. eGenes per eVariant | 3.11 | 1.88 | 3.26 |
| Mean No. Tissues per eVariant | 4.90 | 2.98 | 4.38 |
| Mean No. Tissues per eGene | 9.17 | 9.19 | 9.62 |
| Mean No. eVariants per eGene | 93.77 | 33.17 | 181.16 |

**Supplementary Table 2:** Descriptive Statistics for Enhancer-to-Target Gene Annotation Data Sets

|  | RoadmapLinks | JEME | GeneHancer |
| --- | --- | --- | --- |
| Total No. Regulatory Elements | 1,285,355 | 235,242 | 209,691 |
| Total No. Target Genes | 18,931 | 18,357 | 79,782 |
| Total Mbp Spanned | 257.1 | 247.8 | 331.2 |
| Mean No. Genes per Element | 1.3 | 2.7 | 3.4 |
| Mean No. Elements per Gene | 89.2 | 35.1 | 8.8 |

## Supplementary Figures

**Supplementary Figure 1:**
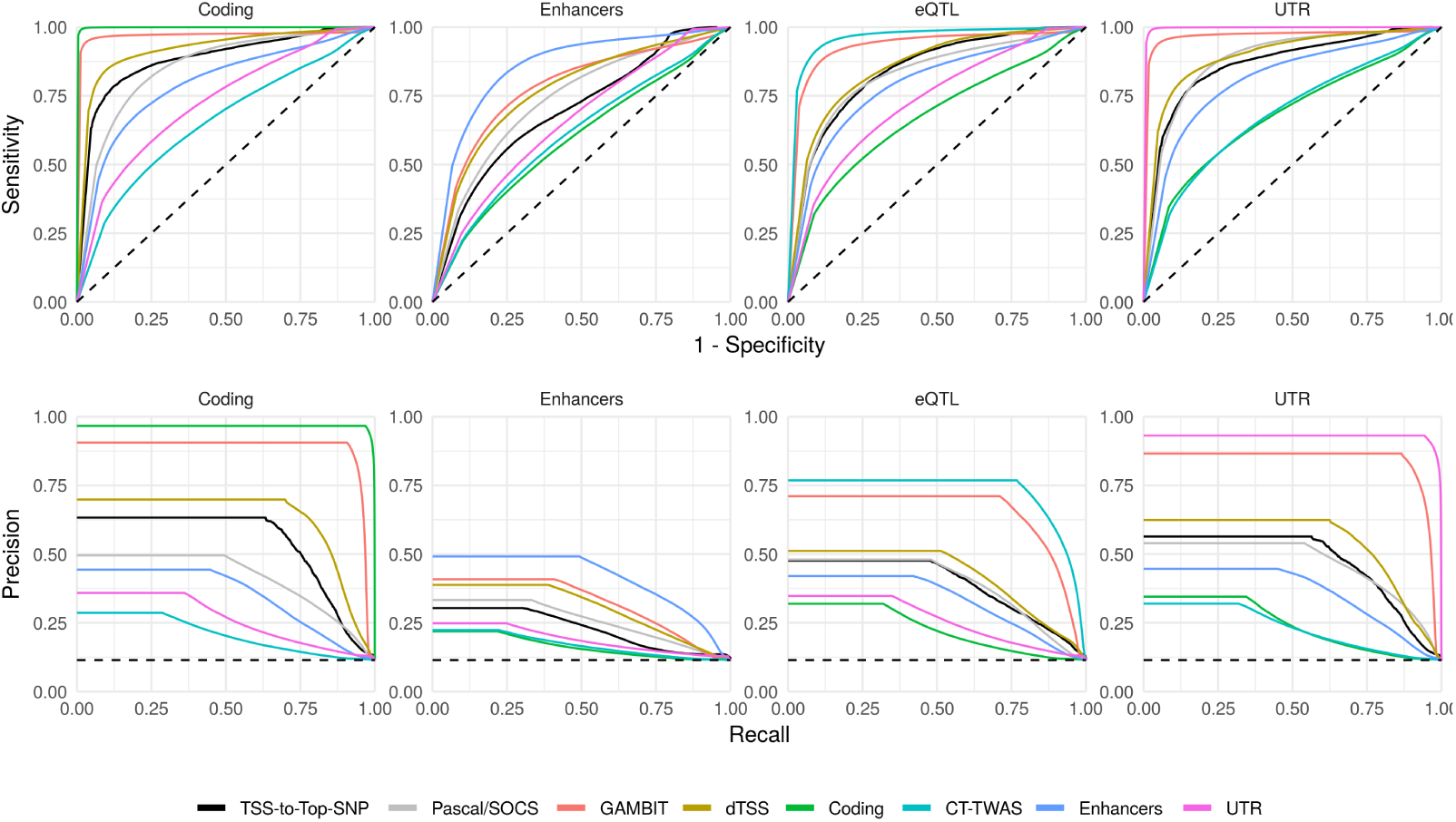
GWAS Simulations: ROC and Precision-Recall Curves. Receiver Operating Characteristic (ROC; top) and Precision-Recall (bottom) curves for each gene-based testing approach (curve color) when either coding, eQTL, enhancer, or UTR variants are causal (plot columns) given locus heritability 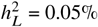. To aggregate results across loci and simulation replicates, we use standardized scores for each method calculated by dividing gene-based scores (e.g., -log_10_-p-values) by the maximum value at the corresponding locus within each replicate. This procedure ensures that curves reflect performance ranking genes at each locus individually. We obtained similar results using the quantile rank of gene-based scores within each locus for each method rather than dividing by the maximum value.

**Supplementary Figure 2:**
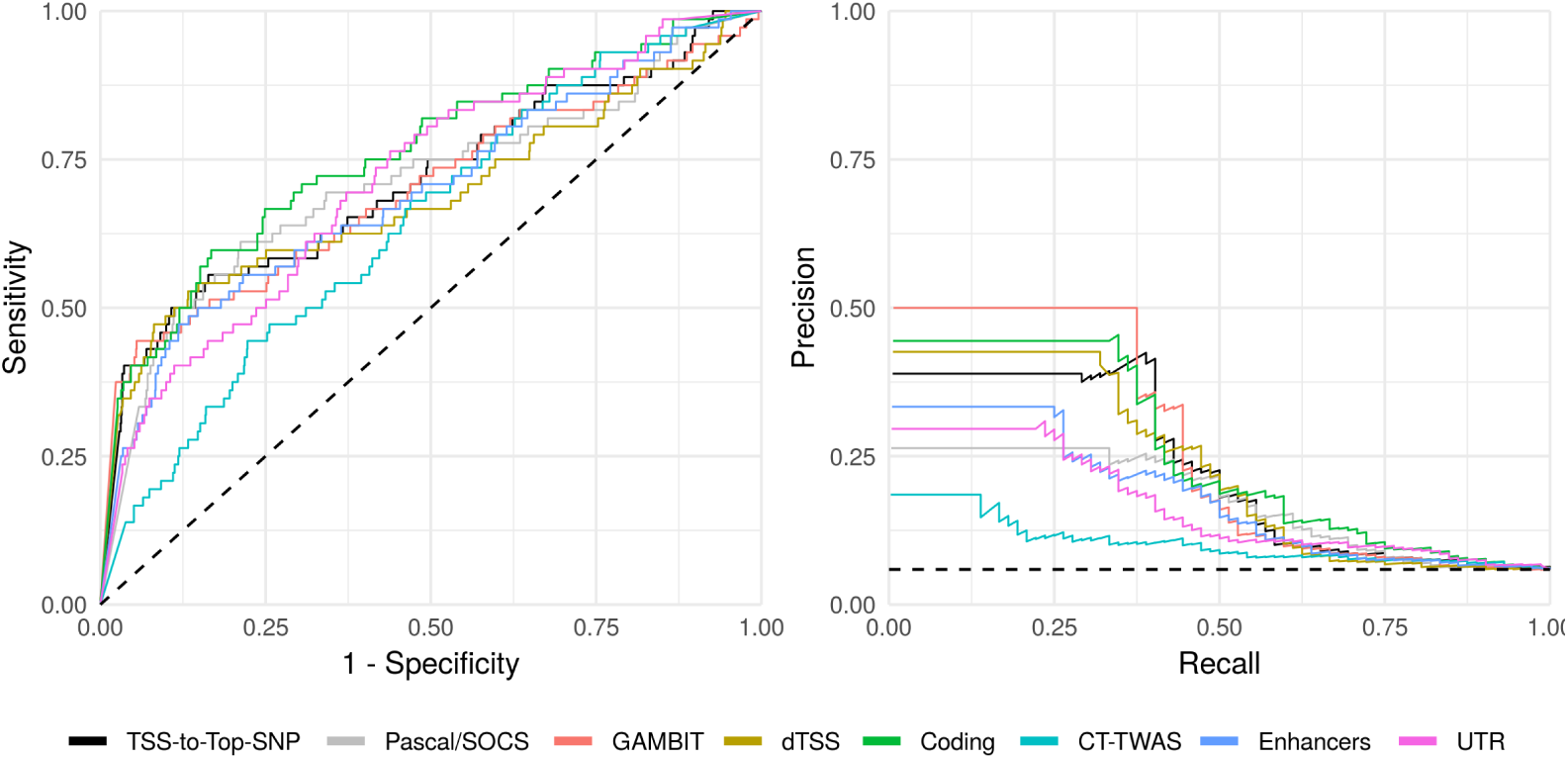
UK Biobank: Sensitivity and Specificity of Gene Ranking Materials and Methods. ROC and Precision-Recall curves for each gene-based association or ranking method across benchmark loci present in both HPO and ClinVar (54 loci in total). To aggregate results across benchmark loci and UK Biobank traits, we use standardized scores for each method calculated by dividing gene-based scores (e.g., -log_10_-p-values) by the maximum value at the corresponding locus. This procedure ensures that curves reflect performance ranking genes at each locus individually. We obtained similar results using the quantile rank of gene-based scores within each locus for each method rather than dividing by the maximum value.

**Supplementary Figure 3:**
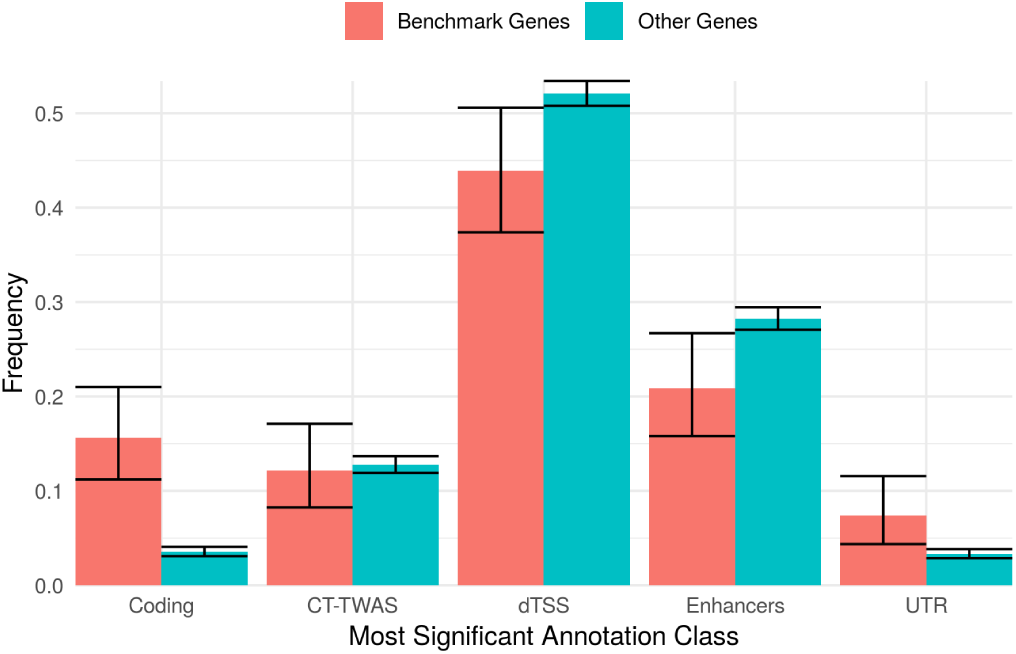
Most Significant Annotation Class for Benchmark vs. Other Genes. Most significant single-annotation test (x-axis) for genes with one or more gene-based p-value ≤5e-6. The proportion of benchmark genes (the union of HPO and ClinVar gene lists) and other genes (not present in either benchmark genes list) for which the indicated annotation class is most significant is shown on the y-axis with 95% confidence intervals. Benchmark genes are strongly enriched for coding associations (odds ratio = 5.03, p-value = 1.3e-16), which is expected due to the selection criteria used to construct benchmark gene lists (described in Materials and Methods).

